# Local adaptation drives the diversification of effectors in the fungal wheat pathogen *Parastagonospora nodorum* in the United States

**DOI:** 10.1101/657007

**Authors:** Jonathan K. Richards, Eva H. Stukenbrock, Jessica Carpenter, Zhaohui Liu, Christina Cowger, Justin D. Faris, Timothy L. Friesen

## Abstract

Filamentous fungi rapidly evolve in response to environmental selection pressures, exemplified by their genomic plasticity. *Parastagonospora nodorum*, a fungal pathogen of wheat and causal agent of septoria nodorum blotch, responds to selection pressure exerted by its host, influencing the gain, loss, or functional diversification of putative effector genes. Whole genome resequencing of 197 *P. nodorum* isolates collected from spring, durum, and winter wheat production regions of the United States enabled the examination of effector diversity and genomic regions under selection specific to geographically discrete populations. A total of 1,026,859 quality SNPs/InDels were identified within the natural population. Implementation of GWAS identified novel loci, as well as *SnToxA* and *SnTox3* as major factors in disease. Genes displaying presence/absence variation and predicted effector genes, as well as genes localized on an accessory chromosome, had significantly higher pN/pS ratios, indicating a greater level of diversifying selection. Population structure analyses indicated two major *P. nodorum* populations corresponding to the Upper Midwest (Population 1) and Southern/Eastern United States (Population 2). Prevalence of *SnToxA* varied greatly between the two populations which correlated with presence of the host sensitivity gene *Tsn1*. Additionally, 12 and 5 candidate effector genes were observed to be diversifying among isolates from Population 1 and Population 2, respectively, but under purifying or neutral selection in the opposite population. Selective sweep analysis revealed 10 and 19 regions of positive selection from Population 1 and Population 2, respectively, with 92 genes underlying population-specific selective sweeps. Also, genes exhibiting presence/absence variation were significantly closer to transposable elements. Taken together, these results indicate that *P. nodorum* is rapidly adapting to distinct selection pressures unique to spring and winter wheat production regions by various routes of genomic diversification, potentially facilitated through transposable element activity.

**Author Summary:** *Parastagonospora nodorum* is an economically important pathogen of wheat, employing proteinaceous effector molecules to cause disease. Recognition of effectors by host susceptibility genes often leads to the elicitation of programmed cell death. However, little is known on the correlation between effector diversity and the spatial distribution of host resistance/susceptibility or the genomic mechanisms of diversification. This research presents the genome resequencing of 197 *P. nodorum* isolates collected from spring, winter, and durum wheat production regions of the United States, enabling the investigation of genome dynamics and evolution. Results illustrate local adaptation to host resistance or susceptibility, as evidenced by population-specific evolution of predicted effector genes and positively selected selective sweeps. Predicted effector genes, genes exhibiting presence/absence variation, and genes residing on an accessory chromosome, were found to be diversifying more rapidly. Additionally, transposable elements were predicted to play a role in the maintenance or elimination of genes. A GWAS approach identified the previously reported *SnToxA* and *SnTox3* as well as novel virulence candidates, as major elicitors of disease on winter wheat. These results highlight the flexibility of the *P. nodorum* genome in response to population-specific selection pressures and illustrates the utility of whole genome resequencing for the identification of putative virulence mechanisms.

## Introduction

Plant pathogenic microorganisms, which are continually in evolutionary conflict with their respective hosts, have developed mechanisms by which they adapt and proliferate. The dynamic nature of fungal genomes, resulting in small and large-scale changes, provides diversification while maintaining essential functions. The prevalence of mobile elements often drives this flexibility, resulting in the creation, abolition, or translocation of genes [1, 2].

Additionally, this activity of mobile elements significantly contributes to the diversification of asexually reproducing fungal pathogens through the rearrangement of chromosomes into lineage specific segments [3, 4]. This phenomenon has recently been described as a compartmentalized ‘two-speed’ genome, consisting of regions of high gene density and equilibrated GC content, as well as a gene-sparse compartment [5,6,7,8]. The regions of low gene density and high repetitive content appear to be hotbeds of rapid evolution, harboring genes encoding virulence determinants known as effectors that manipulate host cellular processes to facilitate infection [7]. However, not all plant pathogenic fungi exhibit such explicit genome architecture but have rapidly evolving genes and transposable elements dispersed evenly throughout the genome, resembling a ‘one-speed’ or ‘one-compartment’ structure [9, 10].

Fungal effector molecules act to modulate the host defense response in various manners, typically dependent on the lifestyle of the pathogen. Pathogen associated molecular patterns (PAMPs), such as the major fungal cell wall constituent chitin, may be recognized by the host and trigger an early defense response known as PAMP triggered immunity (PTI) [11,12,13]. Effectors can successfully subvert this response to permit infection. Ecp6, a fungal effector from *Cladosporium fulvum*, binds chitin and prevents the initiation of PTI [14]. In the context of a biotrophic pathogen, host plants subsequently evolved the means to recognize virulence effectors, triggering a hypersensitive response and sequestration of the pathogen. Recognition is often mediated by host resistance genes in the nucleotide binding leucine-rich repeat (NB-LRR) gene family [15, 16]. Within many biotrophic pathosystems, this interaction between a dominant host resistance gene and a biotrophic effector follows the gene-for-gene model [17]. However, necrotrophic pathogens use effectors to exploit the same host defense cellular machinery. Occurring in an inverse gene-for-gene manner, effectors are recognized by dominant host susceptibility genes resulting in necrotrophic effector triggered susceptibility [18]. The lack of homology or conserved domains between effector proteins hinders efforts towards novel effector discovery. This lack of similarity can be attributed to the rapid evolution in response to local selection pressure exerted by host resistance or susceptibility. However, this can be remedied via thorough genomic and genetic analyses.

Fungi have become a great resource to study genome-wide signatures of selection due to their short generation times and feasibility of whole-genome sequencing. Positive selection of pathogen effector genes, as well as the host resistance/susceptibility genes, drives co-evolutionary processes. The detection of positive selection permits the identification of genes potentially involved in local adaptation to host genotypes or environmental pressures [19]. Examination and comparison of nonsynonymous and synonymous substitution rates within genes has been used to detect evidence of purifying, neutral, or positive selection of putative pathogen virulence genes [20, 21]. More recently, whole-genome sequencing data has been leveraged to identify and compare selective sweep regions between pathogen populations, as well as closely related species. Badouin et al. [22] found a greater prevalence of selective sweeps in *Microbotryum lychnidis-dioicae* compared to its sister species *Microbotryum silenes-dioicae*. Additionally, candidate genes potentially involved in pathogenicity and host adaptation, as well as genes upregulated *in planta* were identified within selective sweep regions. Analysis of a global collection of the barley pathogen *Rhynchosporium commune* identified three major genetic clusters that exhibited unique and generally non-overlapping selective sweep regions hypothesized to stem from ecological variations [23]. Interestingly, genes potentially implicated in response to abiotic stress and not host adaptation were found to be enriched under selective sweep regions. Similarly, evidence for selective sweeps was detected in a worldwide collection of *Zymoseptoria tritici*, with the majority of the identified regions being unique to one of the four postulated populations [24]. However, in this case, genes typically associated with pathogenicity or virulence, such as secreted proteins or cell wall degrading enzymes, were found to underlie selective sweeps.

On a global scale, wheat annually ranks as one of the most widely grown crops and supplies the world with an important source of calories. In 2017, approximately 218.5 million hectares were harvested globally [25]. Common wheat (*Triticum aestivum*), an allohexaploid (AABBDD), is classified by several characteristics including growth habit (spring or winter), bran color (red or white), and hardness of endosperm (soft or hard). Hard red spring wheat and hard red winter wheat are utilized primarily in the bread making industry, whereas flour from soft red winter wheat is often used for cakes and cookies [26]. Durum wheat (*Triticum turgidum* var. *durum*), an allotetraploid (AABB), is primarily used in the pasta industry [26]. Hard red spring wheat and durum wheat are typically grown in the Upper Midwest, hard red winter wheat is typically grown in the Great Plains, and soft red winter wheat cultivation predominates in the southeastern United States [27].

*Parastagonospora nodorum*, a haploid, necrotrophic fungal pathogen of both common wheat and durum wheat, causes significant yield losses and is an annual threat to global wheat production [28]. This pathogen has also emerged as a model organism for the study of plant-necrotrophic specialist interactions. Genomic resources, including the development of near complete reference genomes of diverse isolates bolstered by deep RNA sequencing, have greatly aided the investigation of pathogen virulence. Initially, the *P. nodorum* Australian isolate SN15 was sequenced with Sanger sequencing and subsequently improved via re-sequencing with short-read Illumina technology, RNA-seq data, and protein datasets [29,30,31]. Recently, we developed reference quality genome assemblies of *P. nodorum* isolates LDN03-Sn4 (hereafter referred to as Sn4), Sn2000, and Sn79-1087 using long-read sequencing technology [32]. These telomere to telomere sequences of nearly every *P. nodorum* chromosome, in addition to the annotation of 13,379 genes using RNA-seq, greatly improved the framework for effector discovery. A total of nine effector-host susceptibility factor interactions have been previously identified [18,33,34,35,36,37,38,39,40,41] with three of the pathogen effectors (SnToxA, SnTox1, and SnTox3) having been cloned and validated [35,36,42]. Additionally, the cognate receptor genes of SnToxA and SnTox1, *Tsn1* and *Snn1*, respectively, have also been cloned and characterized in wheat [34, 43]. Novel host sensitivities in winter wheat germplasm have also been identified in response to effectors produced by *P. nodorum* isolates collected in the southeastern United States, which differ from sensitivities and effectors identified in the hard red spring wheat production region of the Upper Midwest [44].

The functionally validated *P. nodorum* effector molecules display typical properties of pathogen effectors, including being small, secreted, cysteine-rich proteins embedded in or adjacent to repetitive regions of the genome. Additionally, these genes exhibit presence/absence variation (PAV), as they are completely missing in avirulent isolates [35,36,42]. This gain or elimination of entire genes, as well as intragenic polymorphisms, is rampant within *P. nodorum* populations, although little correlation between this diversity to the spatial distribution of host susceptibility has been made. Despite the evidence of frequent sexual reproduction in the field, attempts to develop bi-parental sexual populations of *P. nodorum* have been unsuccessful, and the validation of candidate effectors has been accomplished through computational, comparative, and reverse-genetics approaches [35,36,42].

The employment of genome wide association studies (GWAS) overcomes the inherent inability to develop bi-parental fungal populations in *P. nodorum* and allows candidate effectors to be mapped at high-resolution. Due to the widespread availability of genome sequencing, this technique can be relatively easily applied to fungi for effector identification [45]. Gao et al. [46] conducted GWAS in *P. nodorum* using restriction-associated DNA genotyping-by-sequencing (RAD-GBS) data from 191 isolates. Significant marker-trait associations (MTAs) were identified corresponding to effector genes *SnToxA* and *SnTox3*, illustrating the utility of this approach. Additionally, it was determined that linkage disequilibrium (LD) decayed rapidly in *P. nodorum*, highlighting the necessity of higher marker density for the successful identification of causal or linked single nucleotide polymorphisms (SNPs) [46]. GWAS was also used in another pathogen of wheat, *Zymoseptoria tritici*. Sequencing of 103 isolates and subsequent identification of 584,171 single nucleotide polymorphisms (SNPs) enabled the successfully identification of *AvrStb6*, the gene conferring avirulence on wheat lines carrying a functional *Stb6* resistance gene [47, 48].

As plant pathogens rapidly respond to the selection pressure placed upon them by regionally deployed host resistance/susceptibility genes, we hypothesized that this would result in local adaption of effector repertoires which would be visible as an accumulation of population-specific non-synonymous or loss-of-function mutations and genome-wide signatures of selection. This research presents the whole-genome resequencing of 197 *P. nodorum* isolates collected from various wheat growing regions of the United States, enabling the investigation of population structure and regionally-specific gene diversity, including effector diversification. Selective sweep and genic diversity analyses detected unique selection pressures on local *P. nodorum* populations, resulting in the regionally-specific evolution of genes with a predicted virulence function, including putative effector genes that are hypothesized to have specific host susceptibility targets. Genes associated with presence/absence variation were found to be near repetitive elements, indicating a potential role of transposons in the maintenance or elimination of genic diversity. Additionally, a high level of genotypic diversity enabled robust genome-wide association analyses, as illustrated by *SnToxA* and *SnTox3* being significantly associated with disease on both spring and winter wheat lines. GWAS also detected a cell wall degrading enzyme and an entire gene cluster as novel candidates for virulence on winter wheat, laying the foundation for further dissection of this molecular pathosystem.

## Results

### Whole Genome Sequencing and Variant Identification

To quantify and compare patterns of genetic variation in populations of *P. nodorum*, we sequenced full genomes of 197 isolates collected from spring, winter, and durum wheat production regions of the United States. Total sequence per isolate ranged from approximately 126 Mb to 3.01 Gb with a median value of 1.06 Gb. This corresponds to an approximate genome coverage (Sn4 genome size of 37.7 Mb) ranging from 3.34× to 79.8× with a median coverage of 28.08× (S1 Table).

Following filtering for genotype quality and read depth, 1,026,859 SNPs and insertions/deletions (InDels) were identified corresponding to a nucleotide diversity of 0.0062. Analysis of the functional effects of SNPs/InDels revealed a total of 226,803 synonymous and 160,159 non-synonymous polymorphisms within 13,238 genes, with 141 genes lacking any polymorphism (Table 1). Additionally, a total of 5,110 loss of function (LOF) mutations were detected, abolishing the function of 2,848 genes. These variants include 1,464 frameshift mutations, 2,815 premature stop codons, 457 losses of a stop codon, and 374 losses of a start codon (Table 1). The total SNP data set (including intergenic SNPs/InDels) was further filtered for a minor allele frequency of 5% and maximum missing data per marker of 30%, resulting in the identification of 322,613 SNPs/InDels to be used in association mapping analyses.

*P. nodorum* isolates missing genotype calls in greater than 50% of SNP/InDel sites and/or having average gene coverage of less than 95% were discarded from all coverage-based analyses, resulting in a final dataset of 175 *P. nodorum* isolates. Coverage analysis across the 13,379 annotated *P. nodorum* isolate Sn4 gene set identified 882 genes that were deleted in at least one isolate. Individual isolates were annotated as harboring between 5 and 343 gene losses which are distributed throughout the genome. Among genes exhibiting PAV, 70 genes encoded proteins with predicted secretion signals, including the previously characterized *SnToxA*, *SnTox1*, and *SnTox3* [35,36,42]. Overall, no significant differences were observed in the frequency of PAV between predicted effectors, secreted non-effectors, or non-secreted proteins, with PAV frequency rates of 7.8%, 5.2%, and 6.7%, respectively (Pairwise comparison of proportions, FDR adjusted p > 0.21 for all comparisons).

**Table 1.** Functional variants identified in a population of 197 *P. nodorum* isolates

| Variant Category | Total | Population 1 | Population 2 |
| --- | --- | --- | --- |
| Synonymous SNP | 226,803 | 194,313 | 151,965 |
| Non-synonymous SNP | 160,159 | 127,006 | 97,361 |
| Frameshift | 1,464 | 992 | 798 |
| Gain of Stop Codon | 2,815 | 2030 | 1406 |
| Loss of Stop Codon | 457 | 389 | 282 |
| Loss of Start Codon | 374 | 298 | 216 |

### P. nodorum exhibits strong population structure in North America

We next set out to assess if the population structure of *P. nodorum* reflects adaptation to different host populations or geographical distribution. Using a genotypic dataset consisting of approximately one SNP per kb across the entire genome to mitigate potential marker pairs in linkage disequilibrium, STRUCTURE analysis revealed an optimal number of two subpopulations (S1 Figure). All isolates collected from North Dakota, Minnesota, and South Dakota formed a cluster, here termed Population 1 (S1 Figure). Isolates collected from Arkansas, Georgia, Maryland, New York, North Carolina, Ohio, Oregon, South Carolina, Tennessee, Texas, and Virginia formed a second cluster, here termed Population 2 (S1 Figure). Based on the 0.85 threshold of membership probability used to assign isolates to subpopulations, all 17 Oklahoma isolates were not clearly assigned to a cluster. To test the hypothesis that the Oklahoma isolates were admixed between the two major populations, a three-population test was conducted [49]. The f_3_ statistic and z-value were 16.50 and 4.68, respectively. Both values being non-negative indicated a lack of evidence for admixture. This structure and admixture analysis clearly separates isolates based on geographical location rather than wheat cultivar and indicates isolates collected from Oklahoma are likely not admixed, but rather stem from a lineage of the major populations.

Principal components analysis (PCA) revealed similar results to those obtained by Bayesian clustering. The first two principal components, accounting for approximately 12.9% of the cumulative variation, separate Population 1, Population 2, and the isolates from Oklahoma. However, this analysis also indicated a separate cluster corresponding to eight isolates collected from Oregon (lower right of the plot), which likely represent a small subpopulation (Fig. 1). However, due to a comparatively lower population size (n=8), these isolates were removed from any subsequent comparisons between populations due to lack of proper representation (Fig. 1).

**Figure 1.**
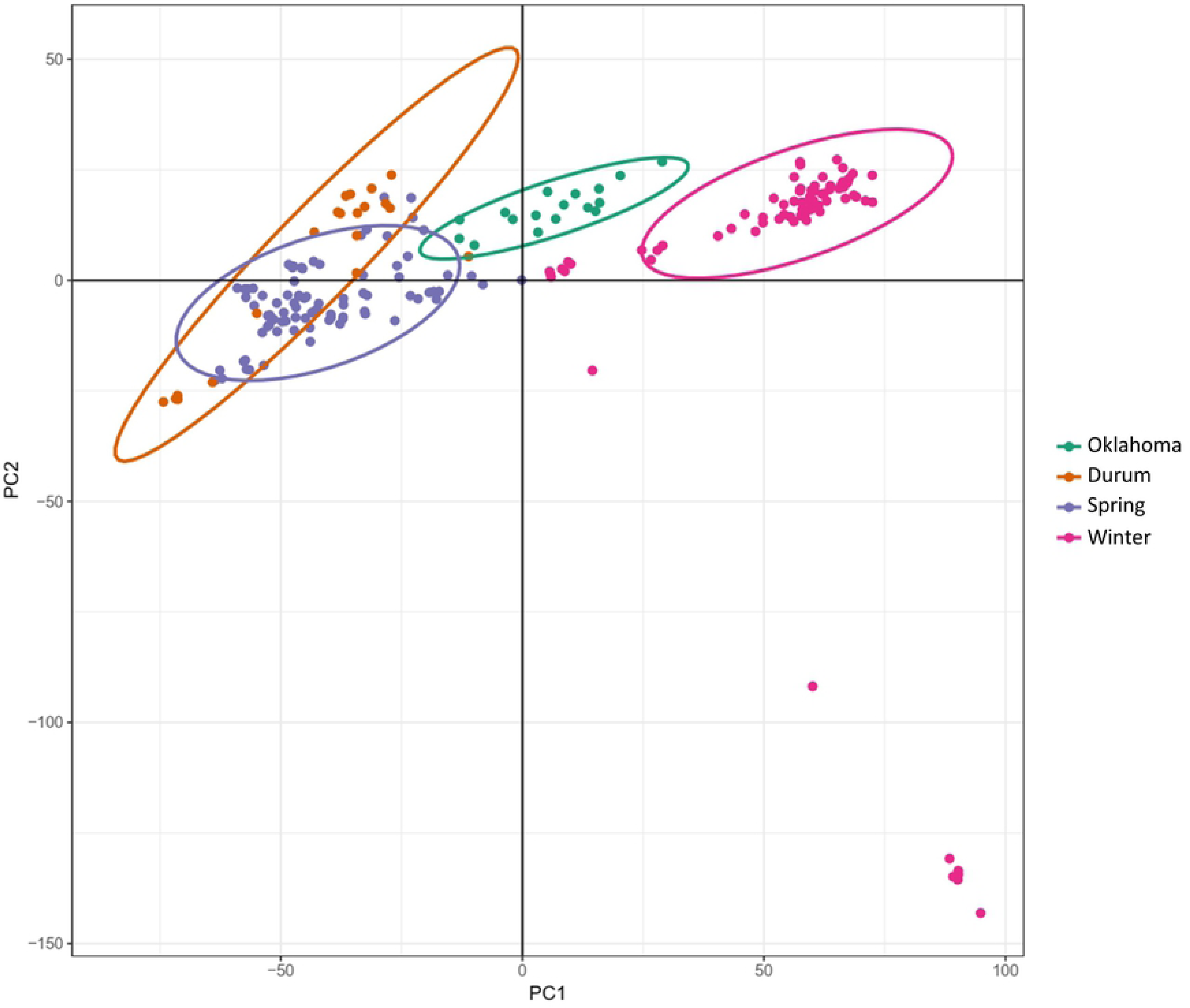
PCA using genotypic data of 50,000 randomly selected markers. PC1 is represented on the x-axis and PC2 is represented on the y-axis. Colored dots correspond to individual isolates and the color signifies the predominant wheat class (or state) of the region from which the isolate was collected. The isolates collected in Oregon are depicted in the lower right of the figure. The color legend is displayed on the right side of the figure.

### High genetic variation in P. nodorum populations

We used our genome-wide inference of genetic variation to compare patterns of genetic diversity and polymorphism distribution in the *P. nodorum* population. Due to the larger number of SNB susceptibility targets that have been validated in spring wheat relative to winter wheat [18], we hypothesized that *P. nodorum* populations specific to spring wheat regions would contain a higher level of diversity and be under balancing selection to maintain a larger virulence gene repertoire. To assess the extent of population differentiation, we computed the parameter F_st_ between each population. The F_st_ statistic was 0.181 when comparing Population 1 to Population 2, indicating moderate differentiation has occurred between isolates collected from each region. To assess the distribution of nucleotide diversity in each *P. nodorum* population, we computed nucleotide variation (π), Watterson’s theta (Θ_W_), and Tajima’s D for each population (Table 2; S2 Figure). Overall, Population 1 had a higher level of nucleotide diversity (0.0059) compared to Population 2 (0.0041). Population 1 also had a greater proportion of segregating sites compared to Population 2, as evidenced by Θ_W_ values of 0.0055 and 0.0037, respectively. Additionally, differences were observed in the estimates of Tajima’s D between the two. Both groups had positive Tajima’s D values caused by an excess of alleles of intermediate frequencies and absence of rare alleles. Population 1 had the highest genome-wide Tajima’s D value at 0.7402. Population 2 had lower, yet still positive, Tajima’s D values of 0.5660. Taken together, these results indicate that isolates comprising Population 1 have a greater level of nucleotide diversity at the genome scale. Also, the positive values of Tajima’s D indicate a possible population contraction or that balancing selection is occurring.

**Table 2.** Population genomics parameters of 172 P. nodorum isolates by population

| Population | $\pi^1$ | $\Theta_W^2$ | Tajima's D<sup>3</sup> |
| --- | --- | --- | --- |
| Population 1 | 0.0059 | 0.0054 | 0.8641 |
| Population 2 | 0.0041 | 0.0037 | 0.5660 |

<sup>1</sup>Nucleotide diversity calculated across entire genome
<sup>2</sup>Watterson's estimator calculated across entire genome
<sup>3</sup>Average Tajima's D value

### Signatures of selective sweeps indicate recently acquired advantageous mutations

Examination of genomic regions having undergone selective sweeps sheds light onto potentially beneficial genes being selected for, as well as facilitates the comparison of selective forces acting on different populations. We hypothesize that due to regional differences in wheat genotypes grown, as well as differing environmental cues, positive selection is acting on different regions of the *P. nodorum* genome within each population. Selective sweep analysis using SweeD revealed 42 and 46 regions having undergone selective sweeps in isolates from Population 1 and Population 2, respectively (Table 3; Fig. 2A). To add further evidence of selective sweeps, predicted sweep regions were compared to population-specific Tajima’s D values calculated in intervals across the genome. This analysis identified a total of 10 and 19 regions in Population 1 and Population 2, respectively, which were predicted as selective sweeps by SweeD and were located in a genomic region with a negative value of Tajima’s D. Interestingly, no genes underlying selective sweep regions were common to both populations, indicating that different selection pressures, likely from regional wheat genotypes or the environment, are being exerted on isolates from each population for the maintenance of beneficial genes.

**Figure 2.**
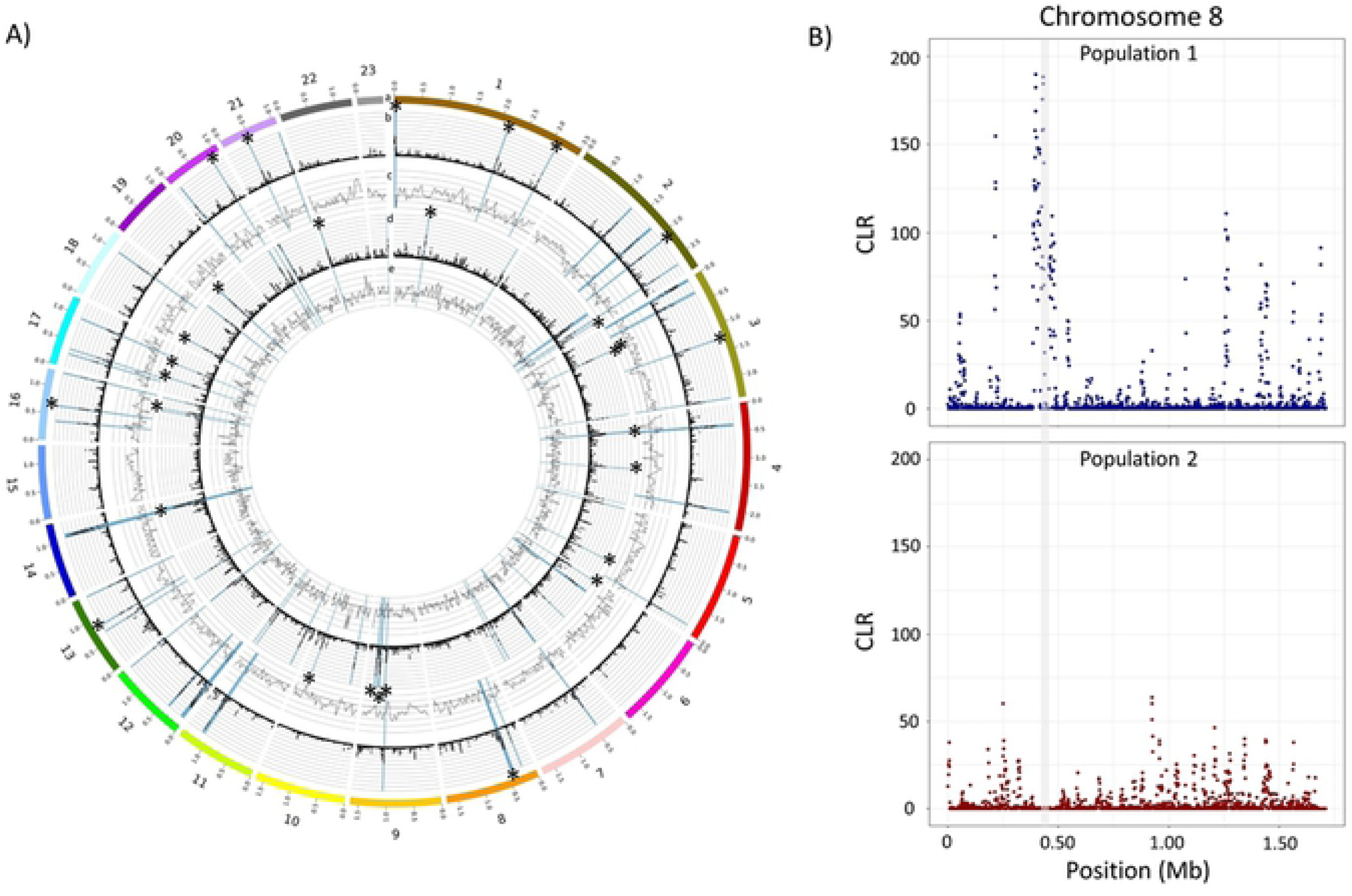
A) Plot illustrating the detected selective sweep loci from both *P. nodorum* subpopulations and sliding window Tajima’s D analysis. ‘*’ signifies a selective sweep region that was detected using SweeD and has a negative average value of Tajima’s D. a) Chromosomes with sizes listed in Mb in 0.5 Mb increments b) Selective sweeps detected in Population 1. Genomic position is displayed on the x-axis. Likelihood values are displayed on the y-axis. Y-axis scale ranges from 0 to 400. Individual dots represent the likelihood value of a single 1 kb window. Highlighted blue regions are the 99^th^ percentile of likelihood values. c) Tajima’s D values of isolates in Population 1 in 50 kb windows in 25 kb steps. Genomic positions are displayed on the x-axis. Tajima’s D values are displayed on the y-axis. The y-axis ranges from −2 to 4. The bold horizontal axis line is 0. d) Selective sweeps detected in Population 2. Genomic position is displayed on the x-axis. Likelihood values are displayed on the y-axis. Y-axis scale ranges from 0 to 400. Individual dots represent the likelihood value of a single 1 kb window. Highlighted blue regions are the 99^th^ percentile of likelihood values. e) Tajima’s D values of isolates in Population 2 in 50 kb windows in 25 kb steps. Genomic positions are displayed on the x-axis. Tajima’s D values are displayed on the y-axis. The y-axis ranges from −2 to 4. The bold horizontal axis line is 0. B) Selective sweep analysis of *P. nodorum* chromosome 8 for Population 1 (top panel) and Population 2 (bottom panel). Position along the chromosome in megabases (Mb) is displayed on the x-axis. The composite likelihood ratio (CLR) is shown on the y-axis. The genomic region harboring *SnToxA* is highlighted in light grey.

**Table 3.** Detection of selective sweeps in two *P. nodorum* populations

| Population | Sweep Regions | Median (kb) | Maximum (kb) | Genes | Genome Coverage (%) |
| --- | --- | --- | --- | --- | --- |
| Population 1 | 10 | 4.5 | 25.0 | 16 | 0.17 |
| Population 2 | 19 | 9.0 | 92.0 | 76 | 1.02 |

Sweep regions detected in Population 1 isolates had a median length of 4.5 kb with the largest region being 25.0 kb. Underlying these regions are 16 genes, of which, 7 are predicted hypothetical proteins with no known functional domains. A total of four genes encode predicted secreted proteins, including one predicted effector. Additionally, five genes (31.3% of genes underlying sweeps) exhibited PAV, which is significantly greater than expected (one gene expected; Fisher’s Exact Test p=0.003). Additionally, genes affected by selective sweeps in Population 1 have a median distance of 6.4 kb to the nearest repetitive element, which although not statistically significant, is substantially closer than the median distance of genes not under selective sweep regions of 13.0 kb (Kruskal-Wallis rank sum test; p = 0.056). Gene ontology enrichment analysis did not identify any significantly overrepresented gene functions, likely due to the low number of genes underlying predicted selective sweep in Population 1 (File S1). A significant selective sweep region was detected on chromosome 8, flanking the *SnToxA* locus, but was not detected in Population 2 (Fig. 2B). This reinforces the hypothesis that *SnToxA* was heavily selected for within the Midwestern population due to the prevalence of *Tsn1*, but was then lost in Population 2 due to the lack of the *Tsn1* gene in the popular local winter wheat cultivars [50].

Analysis of isolates from Population 2 identified 19 selective sweep regions having a median length of 9.0 kb, maximum length of 92.0 kb, and a cumulative length of 385.4 kb, covering 1.02% of the genome (Table 3; Figure 2). The quantity and cumulative size of the predicted sweep regions are larger than those identified in Population 1, indicating that these may be more recent selection events. A total of 76 genes underlie the sweeps detected in Population 2, including 11 genes encoding predicted secreted proteins, one of which is a predicted effector protein. A total of 7 genes (9.2% of genes underlying sweeps) exhibited PAV, which is not significantly greater than expected (five genes expected; Fisher’s Exact test p > 0.05). Additionally, genes underlying selective sweeps in Population 2 had a median distance to the nearest repetitive element of 10.1 kb, which is not significantly closer than genes outside of sweep regions which have a median distance of 13.0 kb (Kruskal-Wallis rank sum test; p=0.63). Gene ontology enrichment analysis did not identify any overrepresented genes underlying selective sweeps specific to Population 2 (S1 File).

### P. nodorum populations harbor different alleles and prevalence of SnToxA, SnTox1, and SnTox3

We then wanted to determine if host sensitivity conferred by a previously characterized effector had influenced *P. nodorum* population structure within the United States. Sensitivity to effectors SnToxA, SnTox1, and SnTox3 conferred by host genes *Tsn1*, *Snn1*, and *Snn3*, respectively, offer a distinct advantage to the pathogen through the strong induction of programmed cell death. The predominant presence of a host sensitivity gene, including novel genes or interactions still undiscovered, may provide a strong selective force towards the maintenance of a given effector, and therefore influence population structure. Coverage analysis was used to determine the presence or absence of the three previously characterized *P. nodorum* effectors *SnToxA, SnTox1*, and *SnTox3* from the natural population (isolates with sufficient coverage, n = 175). Overall, *SnToxA*, *SnTox1* and *SnTox3* were absent from 36.6%, 4.6%, and 41.1%, respectively, from the natural population (Table 4). Prevalence of effector genes was also examined by subpopulation. The clear majority of isolates retained a functional *SnTox1* gene, as 0% and 12.5% of isolates from Population 1 and Population 2, respectively, harbored *SnTox1* gene deletions. *SnTox3* was observed to be absent from 38.3% and 48.4% of isolates from Population 1 and Population 2, respectively. A stark difference was observed when comparing the presence of *SnToxA* between the two populations. Within Population 1, containing isolates from the Upper Midwest, only 4.3% of isolates lacked *SnToxA*. However, among Population 2, isolates from the Southern, Eastern, and Pacific Northwest, 93.8% lacked *SnToxA* (Table 4). Additionally, *SnToxA* was present in 100% (n = 17) of the isolates collected from Oklahoma, which were not placed into a major population. These results indicate a strong selection pressure has been placed on maintaining *SnTox1* in the entire natural populations likely due to its dual function [42]. Additionally, the presence of *SnToxA* has been selected for in Population 1, but on the other hand selected against in Population 2, likely due to regional differences in deployment of the host sensitivity gene *Tsn1*.

**Table 4.** Functional variants identified in *SnToxA, SnTox1, and SnTox3*

| Variant Category | *SnToxA* | *SnTox1* | *SnTox3* |
| --- | --- | --- | --- |
| Coding Sequence (bp) | 534 | 351 | 693 |
| Synonymous SNP | 8 | 0 | 3 |
| Non-synonymous SNP | 3 | 9 | 5 |
| Gain of Stop Codon | 1 | 0 | 0 |
| Loss of Start Codon | 0 | 0 | 1 |
| Deletion (%) Population 1 | 4.3 | 0 | 38.3 |
| Deletion (%) Population 2 | 93.8 | 12.5 | 48.4 |
| Overall Deletion (%) | 36.6 | 4.6 | 41.1 |

In addition to the PAV exhibited by all three effectors, functional diversity was also detected within the coding regions of each gene. Throughout the entire population, *SnToxA* harbored 12 mutations, including eight synonymous SNPs, three nonsynonymous SNPs, and one SNP introducing a premature stop codon (Table 4). A total of four unique nucleotide haplotypes and protein isoforms were detected. The SNP inducing a premature stop codon was found in a single isolate collected on winter wheat from Ohio. Within the *SnTox1* coding region, no synonymous changes were detected, however, nine nonsynonymous SNPs were identified (Table 4). The nine non-synonymous changes form nine unique nucleotide haplotypes and protein isoforms. *SnTox3* was found to contain a total of nine SNPs, including three synonymous changes, five nonsynonymous SNPs, and one SNP causing the loss of the start codon (Table 4). The variants detected within *SnTox3* collapse into five nucleotide haplotypes and four protein isoforms. The start codon mutation was only observed in one isolate, collected on durum wheat in western North Dakota.

### Disease Phenotyping

Previous research has identified the presence of novel necrotrophic effectors in *P. nodorum* isolates collected from the southeastern United States and cognate host sensitivities specific to winter wheat germplasm [44]. In order to better characterize disease reactions of winter wheat to a diverse pathogen collection and potentially identify genes contributing to virulence via GWAS, wheat lines Alsen (*Tsn1* control), Jerry (hard red winter), TAM105 (hard red winter), ITMI38 (*Snn3* control), Massey (soft red winter), and F/G95195 (soft red winter) were inoculated with 197 *P. nodorum* isolates collected from spring, winter, and durum wheat production regions of the United States. Average disease reactions on spring wheat line Alsen ranged from 0.25 to 5.00 with an average of 3.34 (Fig. 3). Disease reaction on hard red winter wheat line Jerry ranged from 0.25 to 4.25 with an average of 2.93 (Fig. 3). Disease reaction on hard red winter wheat line TAM105 ranged from 0.13 to 4.13 with an average of 2.75 (Fig. 3). An increased disease reaction correlated with the presence of a functional *SnToxA* gene, indicating that the SnToxA-Tsn1 interaction is largely responsible for disease on these wheat lines. Disease reaction on the recombinant inbred wheat line ITMI38 ranged from 0 to 3.50 with an average of 1.49 (Fig. 3). Disease reaction on soft red winter wheat line Massey ranged from 0.00 to 2.00 with an average of 0.72 (Fig. 3). Disease reaction on soft red winter wheat line F/G95195 ranged from 0.00 to 3.875 with an average of 1.51 (Fig. 3). Disease reaction correlated with the presence of *SnTox3* which indicated that the SnTox3-Snn3 interaction is the main facilitator of disease on these wheat lines. Interestingly, the disease reactions on Massey were comparatively lower, indicating that although SnTox3 is an effective virulence factor, an underlying host resistance not present in the other wheat lines may exist.

**Figure 3.**
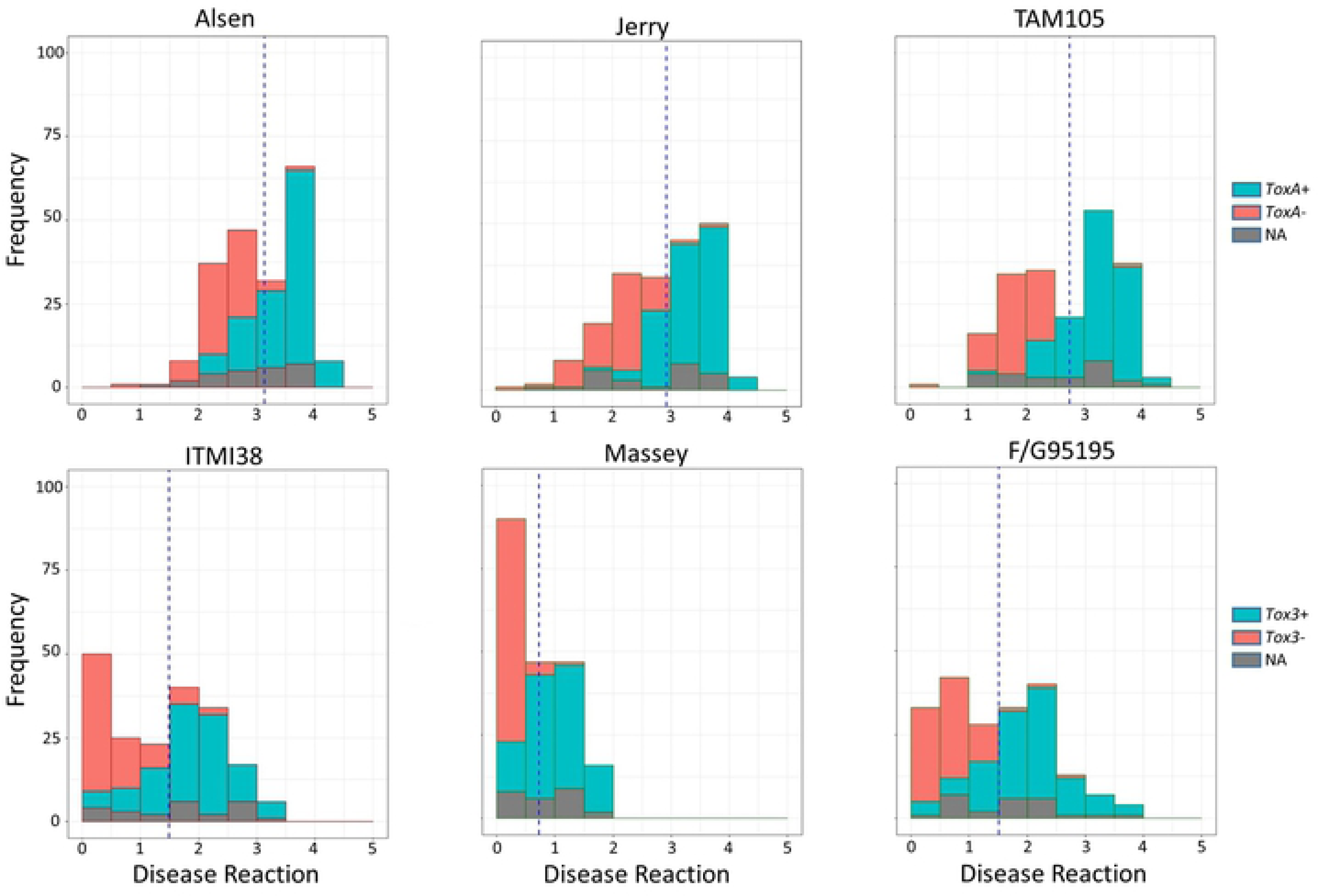
Histograms depicting the distribution of disease reactions to 197 *P. nodorum* isolates on wheat lines Alsen (*Tsn1*), Jerry, TAM105, ITMI38 (*Snn3*), Massey, and F/G95195. Disease reactions are displayed in bins along the x-axis and frequency is illustrated on the y-axis. Bars are shaded to illustrate the proportion of presence/absence of the corresponding effector (*SnToxA* or *SnTox3*) or missing data (NA) per bin. Light blue corresponds to the presence of the effector, dark pink corresponds to the absence of the effector, and grey corresponds to missing data.

### GWAS provides new virulence enhancement candidates and insight into local levels of linkage disequilibrium

Previous studies have identified effectors that interact with host susceptibility genes derived from spring wheat germplasm. Novel effector-susceptibility gene interactions have been identified that are specific to the winter wheat gene pool [44]. To identify candidate virulence genes underlying these novel interactions on winter wheat lines, as well as determine if previously identified effectors play a role in disease development, phenotypic and genotypic data were utilized to conduct a GWAS. Further filtering the quality SNPs/InDels identified from the *P. nodorum* natural population (n = 197) for a minor allele frequency of 5%, a total of 322,613 markers were used to identify associations with virulent phenotypes. Using a mixed linear model incorporating a kinship matrix, a total of 174 and 277 markers were identified as significant in *P. nodorum* for virulence on winter wheat lines Jerry and TAM105, respectively (Fig. 4A). The same marker approximately 51.2 kb upstream of *SnToxA* on chromosome 8 was detected with the highest significance on both wheat lines and the PAV of *SnToxA* was also highly significant on each line (Fig. 4A). *SnToxA* resides in an approximately 112.1 kb isochore region of chromosome 8 characterized by low GC content. Due to the repetitive nature of this region, SNPs could not be reliably called, leaving the PAV of *SnToxA* as the only marker within this region. Among the 174 markers significantly associated with virulence on winter wheat line Jerry, 54 were in a 62.6 kb genomic region downstream and 111 were in an 87.0 kb region upstream of *SnToxA*. A total of eight markers were identified outside of the *SnToxA* locus, including four markers on chromosome 4, one marker on chromosome 10, two markers on chromosome 11, and one marker on chromosome 14 (S3 Figure). Out of the 277 significant markers identified for virulence on TAM105, 80 were in a 66.2 kb genomic region downstream and 173 were in an 88.1 kb region upstream of *SnToxA*. A total of 23 markers were detected outside of the *SnToxA* locus, including four markers on chromosome 1, one marker on chromosome 2, one marker on chromosome 3, three markers on chromosome 4, eight markers on chromosome 5, three markers on chromosome 9, two markers on chromosome 10, and one marker on chromosome 12 (S3 Figure).

**Figure 4.**
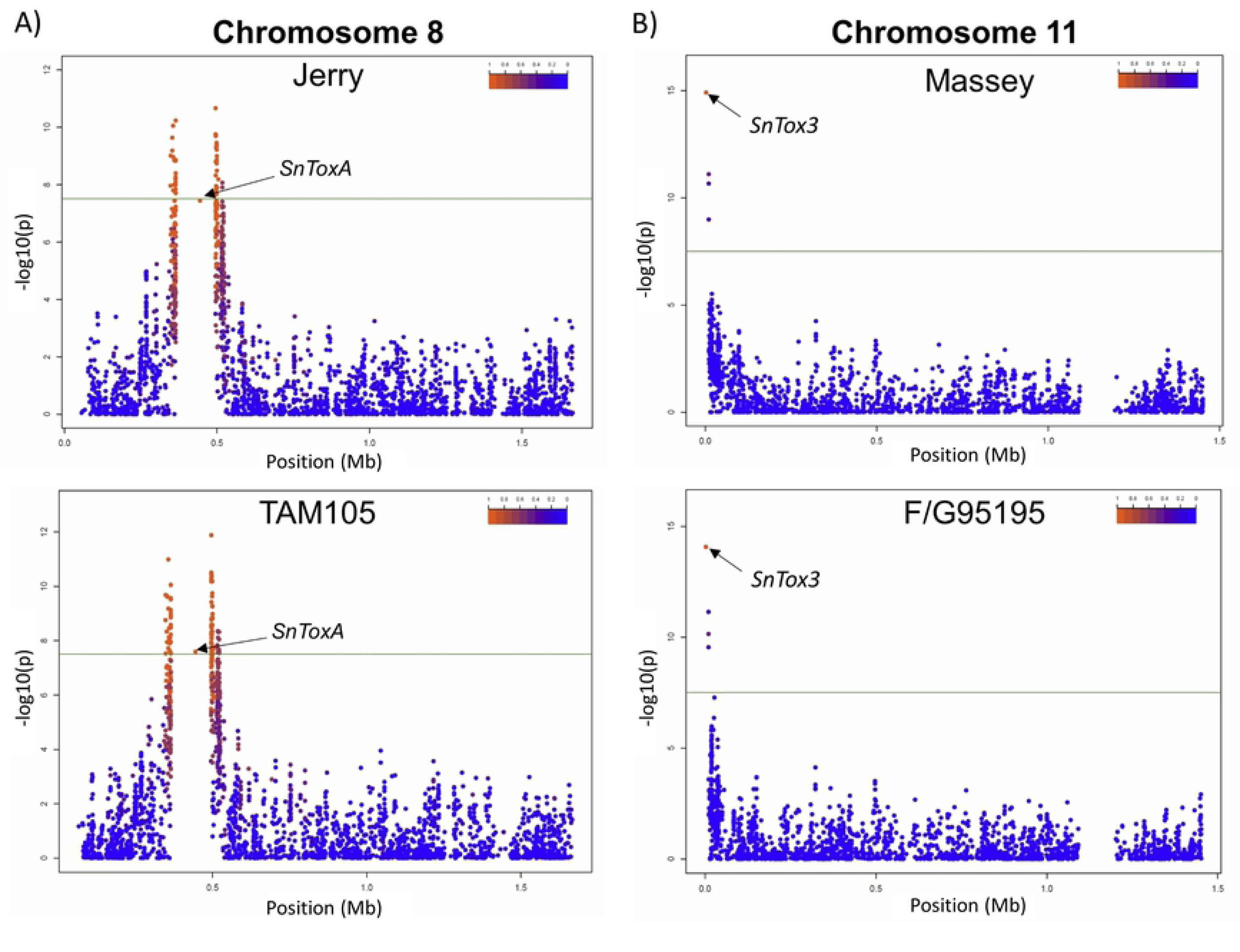
Genome-wide association study (GWAS) detecting significant associations with virulence on wheat lines Jerry, TAM105, Massey, and F/G95195. (A) Manhattan plot of *P. nodorum* chromosome 8 illustrating a highly significant association with the *SnToxA* locus for isolates inoculated onto hard red winter wheat lines Jerry and TAM105. Markers are represented by dots which are colored corresponding to the level of linkage disequilibrium (R^2^) with the most significant marker and are order by position on the x-axis. The significance of each marker expressed as –log10(p) is displayed on the y-axis. (B) Manhattan plot of *P. nodorum* chromosome 11 illustrating a highly significant association with the *SnTox3* locus for isolates inoculated onto soft red winter wheat lines Jerry and TAM105. Markers are represented by dots which are colored corresponding to the level of linkage disequilibrium (R^2^) with the most significant marker and are ordered by position on the x-axis. The significance of each marker expressed as –log10(p) is displayed on the y-axis.

Candidate genes involved in virulence on TAM105 were also identified underlying novel loci. The significant SNP on chromosome 1 at position 2,989,164 bp was within a glycosyl hydrolase family 11 gene, implicated in the degradation of plant cell walls. A high level of polymorphism was detected within this gene, as evidenced by 18 SNPs, including seven non-synonymous SNPs, all within the predicted functional domain. The most significant SNP was a non-synonymous change from valine to isoleucine at amino acid position 116. Directly flanking a significant SNP at position 1,484,536 on chromosome 5 by 1364 bp was an entire gene cluster exhibiting PAV. This cluster was absent from approximately 53% of the isolates and was comprised of a NAD(P)-binding monoxygenase, aldehyde dehydrogenase, aromatic ring opening dioxygenase, acyl esterase/dipeptidyl peptidase, and a transcription factor. In isolates harboring the cluster, the transcription factor was observed to harbor a high level of polymorphism, including 29 non-synonymous and 11 synonymous SNPs.

Interestingly, the PAV of *SnToxA* was not the most significant marker associated with virulence on winter wheat lines Jerry and Massey but still exhibited a high LD with the most significant marker (R^2^=0.97). Additionally, LD extended approximately 48.6 kb downstream and 4.76 kb upstream of *SnToxA* before decaying to R^2^ levels below 0.20, explaining the large number of SNPs identified in association with virulence. These results, confirmed by sensitive reactions of Jerry and TAM105 to infiltrations with SnToxA (Fig. 5A), indicated that SnToxA was the major effector facilitating infection on these two winter wheat lines. However, other candidate genes contributing to disease development in a quantitative manner were identified at significant genomic loci and provide candidate genes for further investigation.

**Figure 5.**
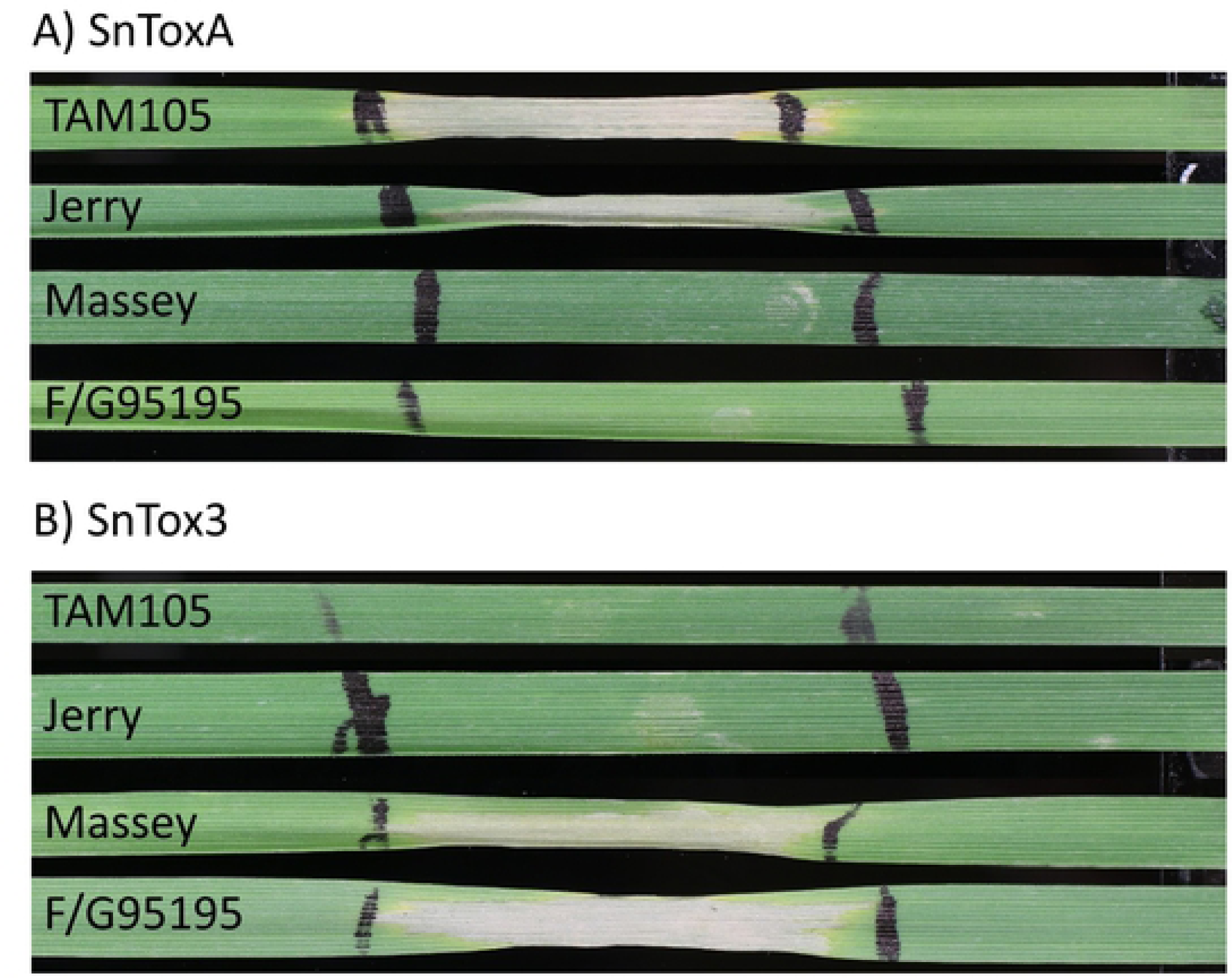
Sensitivity of winter wheat lines TAM105 (hard red), Jerry (hard red), Massey (soft red), and F/G95195 (soft red) to infiltrations with SnToxA (A) and SnTox3 (B). TAM105 and Jerry are sensitive to SnToxA but insensitive to SnTox3, indicating that they possess a functional *Tsn1* and lack *Snn3*. Massey and F/G95195 are insensitive to SnToxA but sensitive to SnTox3, indicating that they lack a functional *Tsn1* but harbor *Snn3*.

Association mapping analyses using a mixed linear model incorporating three principal components (15.9% cumulative variation) as fixed effects and a kinship matrix as a random effect revealed four markers significantly associated with virulence on winter wheat line Massey. The most significant association corresponds to the PAV of *SnTox3* on chromosome 11, with the three remaining significant markers being SNPs located in an approximately 6.0 kb genomic region upstream of the *SnTox3* gene (Fig. 4B). Using the same mixed linear model, a total of ten significant markers were detected in association with virulence on winter wheat line F/G 95195. Similar to virulence on Massey, *SnTox3* PAV was the most significant locus identified, with nine additional significant SNPs located in an ∼23.1 kb upstream region (Fig. 4B). LD decayed to levels below 0.20 approximately 6.5 kb upstream of *SnTox3*. As this gene is located near the telomere, no distal markers were identified, therefore, no LD estimations could be made downstream of *SnTox3*. As no markers outside of the *SnTox3* genomic region were found significant, these results indicate that SnTox3 is the lone major segregating effector governing virulence on the winter wheat lines Massey and F/G 95195, which is confirmed by the sensitivity to infiltrations of SnTox3 (Fig. 5B).

### Gene ontology enrichment analyses show an excess of non-synonymous SNPs within putative virulence genes

For the examination of putative functions of genes undergoing diversification, gene ontology enrichment analysis was conducted. Using 882 genes exhibiting PAV in at least one *P. nodorum* isolate, 16 molecular function gene ontology terms were significant at FDR-adjusted p < 0.10 (S1 File). Genes involved in the binding of nucleotides, small molecules, organic cyclic/heterocyclic compounds, anions, and carbohydrate derivatives were found to be significantly enriched (File S1).

A total of 2,848 genes harbored at least one LOF mutation and were used in gene ontology enrichment analysis, identifying six molecular function ontology terms significant at FDR adjusted p < 0.10 (S1 File). Genes potentially involved in virulence through production or manipulation of reactive oxygen species, as well as genes implicated in molecule binding were enriched, as evidenced by the most significant biological process terms being oxidation-reduction processes, as well as binding of coactors, flavin adenine dinucleotides, heme, tetrapyrrole, and iron ions (File S1).

Examination of gene locations and corresponding pN/pS ratios revealed a significantly higher pN/pS ratio of genes residing on the accessory chromosome (Chromosome 23) compared to the core chromosomes (Pairwise Wilcoxon Rank Sum Test, p < 2 × 10^−16^, w = 333,700). Genes on the accessory chromosome had a median pN/pS value of 0.50 compared to the genome-wide median of 0.20 (Fig. 6). This was also compared to a random subsample of equal size (n=126 genes). The pN/pS medians of the five random subsamples ranged from 0.19-0.22, which is significantly different than the 126 genes residing on the accessory chromosome (Pairwise Wilcoxon Rank Sum Test, p < 2 × 10^−16^). This indicates that genes residing on the accessory chromosome are evolving faster and this genomic compartment may serve as a hotbed of adaptation. A total of 423 and 484 genes from Population 1 and Population 2, respectively, exhibited pN/pS values greater than 1 and were examined for gene ontology enrichment. In Population 1, within the ontology of molecular function, metallopeptidase activity was found significant at p = 0.00061 (0.4 genes expected, 4 genes observed; FDR-adjusted p-value = 0.35). Although not significant following p-value adjustment, the molecular function chitin binding was the most enriched ontology term in Population 2 at p = 0.015 (0.53 genes expected, 3 genes observed).

**Figure 6.**
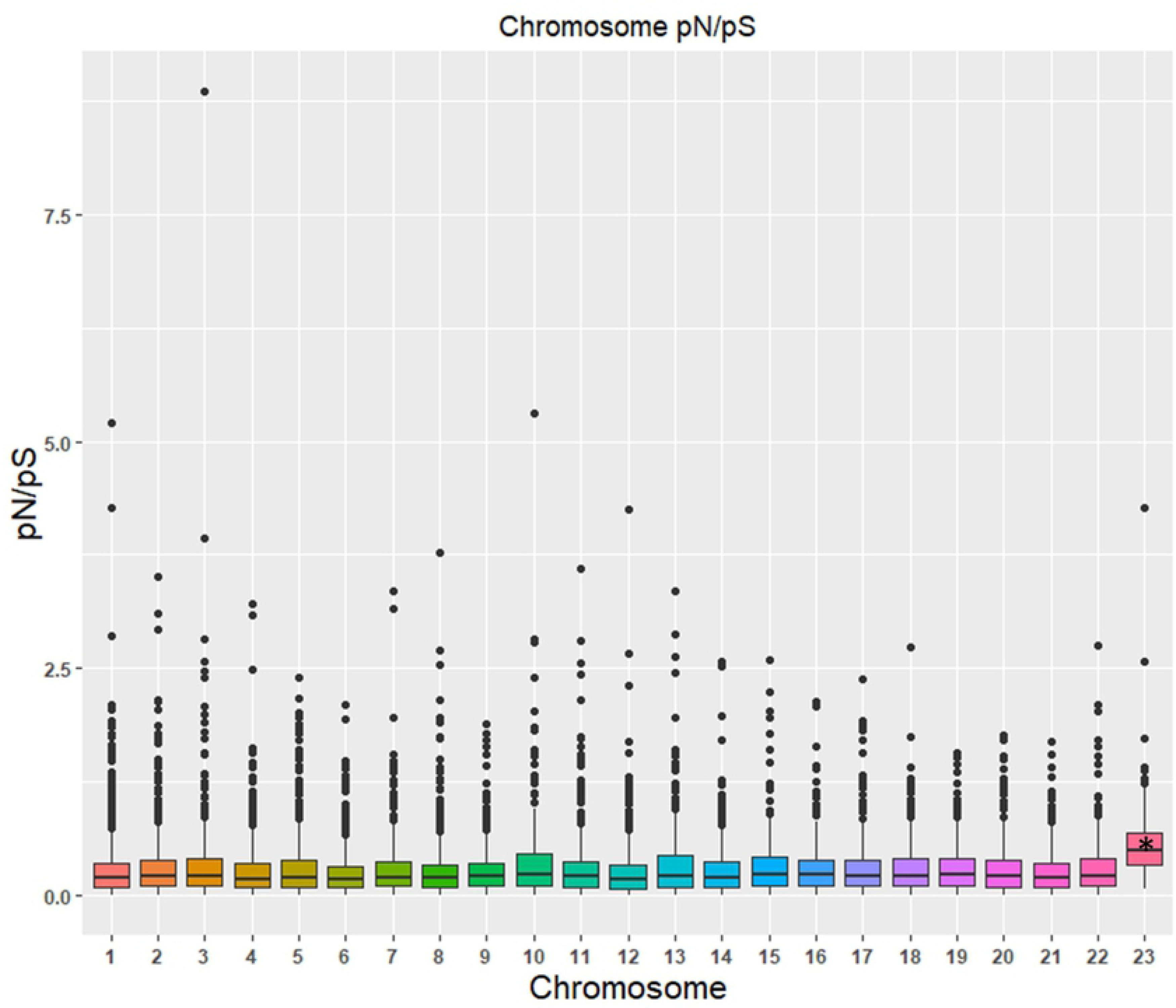
Distribution of pN/pS ratios of genes across all *P. nodorum* chromosomes. Specific chromosomes are displayed on the x-axis and pN/pS ratios are displayed on the y-axis. ‘*’ indicates significantly different than each pairwise comparison at p < 2 × 10^−16^ (Pairwise Wilcoxon Rank Sum test, p < 0.01). Outliers with pN/pS values greater than 10 were omitted.

### Predicted effectors evolve faster than non-effectors

As effectors play an essential role in the development of disease and are directly affected by the selection pressure exerted by host resistance, we next wanted to determine if these genes are preferentially accumulating nonsynonymous changes, as well as if these changes are population specific. Additionally, extreme forms of diversification including the abolition of gene function via loss-of-function mutations or the direct elimination of an effector gene were analyzed. Analysis of the *P. nodorum* isolate Sn4 annotated gene set revealed a total of 1,020 proteins containing a predicted secretion signal and lacking a predicted transmembrane domain. Further analysis using EffectorP revealed 219 of these proteins to be predicted effectors. This candidate effector list was used for further comparative analyses between the derived populations, excluding the isolates from Oklahoma and Oregon. Predicted effector proteins were observed to accumulate a greater number of nonsynonymous SNPs compared to secreted non-effectors or non-secreted proteins (Pairwise Wilcoxon Rank Test, p < 0.01) (Fig. 7A). Genes encoding predicted effectors had average pN/pS ratios of 0.36, whereas secreted non-effectors and secreted proteins had average pN/pS ratios of 0.22 and 0.30, respectively. Also, the clear majority of predicted effector proteins did not have a predicted function, with only 27.6% having predicted functional domains, compared to 64% of secreted non-effector proteins. Only three predicted effector genes lacking a predicted functional domain were completely fixed between the two populations and no synonymous, nonsynonymous, LOF, or PAV mutations were identified. Additionally, 17 genes harbored only nonsynonymous SNPs and therefore, no pN/pS ratios could be calculated. Only two of these predicted effectors contained predicted functions, including chitin and ubiquitin binding. Conversely, 17 predicted effectors had pN/pS ratios of zero due to the presence of only synonymous SNPs. Of these genes lacking nonsynonymous SNPs, 10 contained predicted functional domains, including those involved in cell wall degradation, secretion, and synthesis of phytotoxins. A set of five genes encoding predicted effector proteins, including previously characterized effector *SnTox3*, were found to have pN/pS ratios equal to or less than 1 in Population 2 but appeared to be diversifying in Population 1 (S3 Table). All five proteins lacked predicted functional domains. Also, seven predicted effectors lacking functional domains were found to have only non-synonymous SNPs (and pN/pS could not be calculated) in Population 1, but lacked non-synonymous changes in Population 2 (S3 Table). Conversely, four genes lacked an accumulation of nonsynonymous changes in Population 1 but had pN/pS ratios greater than 1 in Population 2. None of these proteins had predicted functional domains. Additionally, a gene encoding a predicted effector with a chitin binding domain lacked nonsynonymous polymorphism in Population 1 but harbored nonsynonymous SNPs in Population 2 (S3 Table).

**Figure 7.**
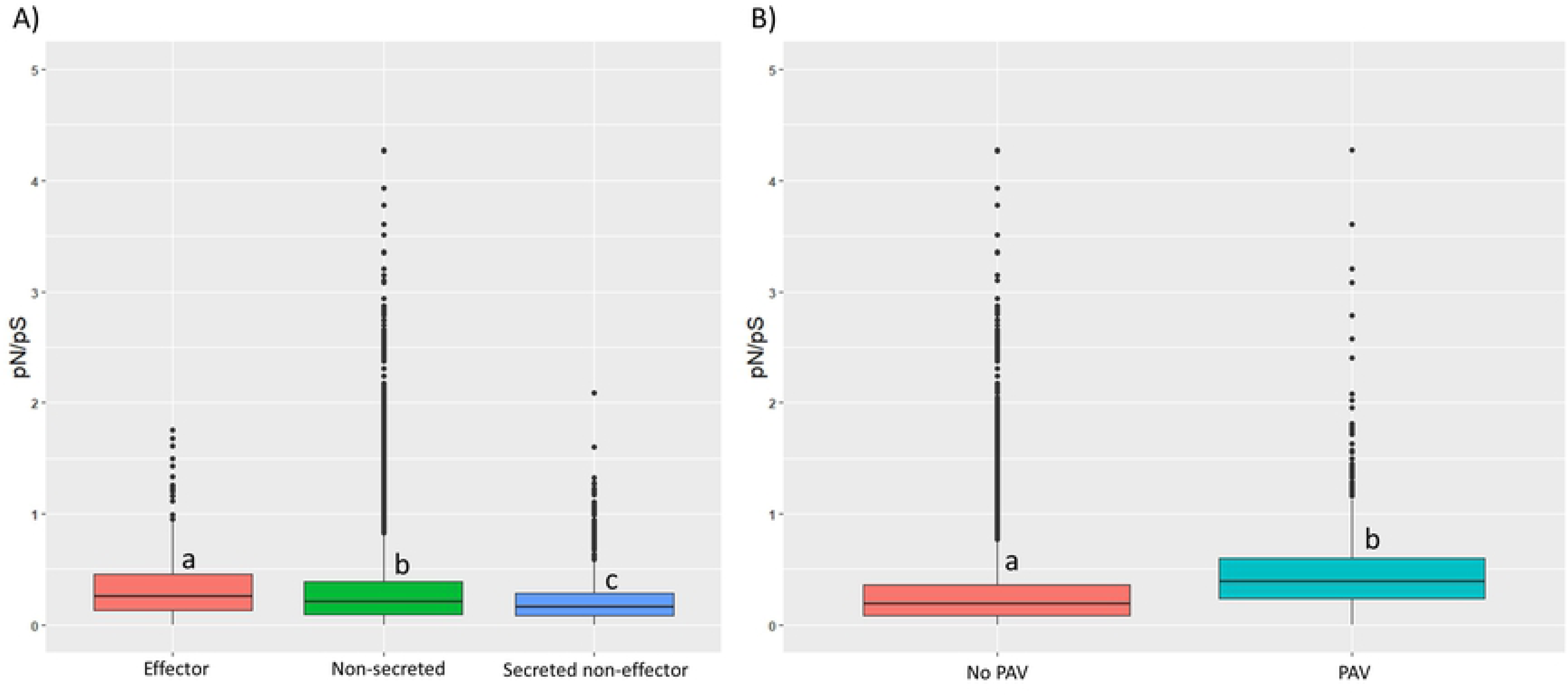
Distribution of pN/pS ratios among genes encoding (A) predicted effectors, secreted non-effectors, non-secreted proteins, and (B) genes exhibiting PAV. Categories are displayed on the x-axes and the log transformed pN/pS ratio is illustrated on the y-axes. Color legends are displayed to the right of each figure. Letter codes correspond to statistically different groups (Pairwise Wilcoxon Rank Test, p < 0.01). Outliers of pN/pS values greater than 10 are not displayed.

LOF mutations affected 45 genes encoding predicted effector proteins. Within Population 1, twelve effector genes harbored LOF mutations but remained intact in all isolates from Population 2. Of these LOF mutations specific to Population 1, eight lacked functional domains, with the remaining genes consisting of a peroxidase, polyketide cyclase, domain of unknown function (1996), and the mutation causing a premature stop codon in SnToxA mentioned previously. Conversely, LOF mutations were identified in 14 effector genes in isolates from Population 2 but remained functional in isolates from Population 1. One gene from this subset encodes a protein with a heterokaryon incompatibility domain, with the remaining proteins lacking functional predictions. A total of 19 genes encoding predicted effector proteins that harbored LOF mutations were common to both populations, all of which are hypothetical proteins except one predicted effector annotated as a blastomyces yeast phase specific protein. Overall, isolates from Population 1 had a significantly higher number of genes affected by LOF mutations, with 2,193 genes having frameshifts, loss of start codons, loss of stop codons, or gain of stop codons, compared to 1,849 detected in isolates from Population 2 (Kruskal-Wallis test, p-value < 0.001).

Differences were also observed with genes exhibiting PAV between populations. As previously mentioned, 70 genes encoding predicted secreted proteins were observed to exhibit PAV, out of which, 17 encoded proteins predicted to be effectors. Two genes, including *SnTox1*, were present in all isolates from Population 1, whereas the PAV was segregating in the isolates from Population 2. Neither of these genes encoded proteins with predicted functional domains although SnTox1 has been shown to bind chitin. A total of four genes exhibited segregating PAV in Population 1 but were present in all isolates from Population 2. Among the proteins encoded by these four genes, two did not have predicted functions and the remaining two consisted of the previously characterized phytotoxic cerato-platanin gene *SnodProt1* [51, 52] and an oxidoreductase. Interestingly, *SnodProt1* was absent only in isolates collected from durum wheat in North Dakota. Additionally, all genes (effector, secreted non-effector, or non-secreted) exhibiting PAV were observed to have significantly higher pN/pS ratios compared to those genes found in all isolates (Kruskal-Wallis test, p < 2 × 10^−16^), indicating that not only are they being strongly selected via elimination or gain of the entire gene, but are diversifying within isolates harboring a functional copy (Fig. 7B).

### Repetitive elements are associated with PAV loci

As repetitive elements have been observed to be a large contributing factor in the dynamic nature of the fungal genome [1, 2], we next wanted to examine the repeat content of *P. nodorum* and its relation to the diversity observed within the natural population. Repeat annotation classified 2,223,841 bp of the Sn4 genome as repetitive content, consisting of LTR/Copia (54.37%), LTR/Gypsy (22.33%), DNA/TcMar-Fot1 (11.64%), LINE/Penelope (2.60%), Satellites (0.29%), and unknown (8.77%) elements, all of which represented approximately 5.90% of the entire genome. Proximity of genes to the nearest repeat element were calculated to determine if transposable or repetitive elements influenced gene diversification. When comparing proximity to repetitive elements of genes exhibiting PAV, a large discrepancy was observed. Genes present in all isolates were a median distance of 13.3 kb away from the nearest repetitive element, while genes exhibiting PAV were significantly closer, being only a median distance of 5.8 kb away (Pairwise Wilcoxon Rank Sum Test, p < 2.0 × 10^−16^; Figure 8). These results indicate that transposable or repetitive element activity may be influencing the gain or loss of genes and are a driving factor in the constantly evolving fungal genome.

**Figure 8.**
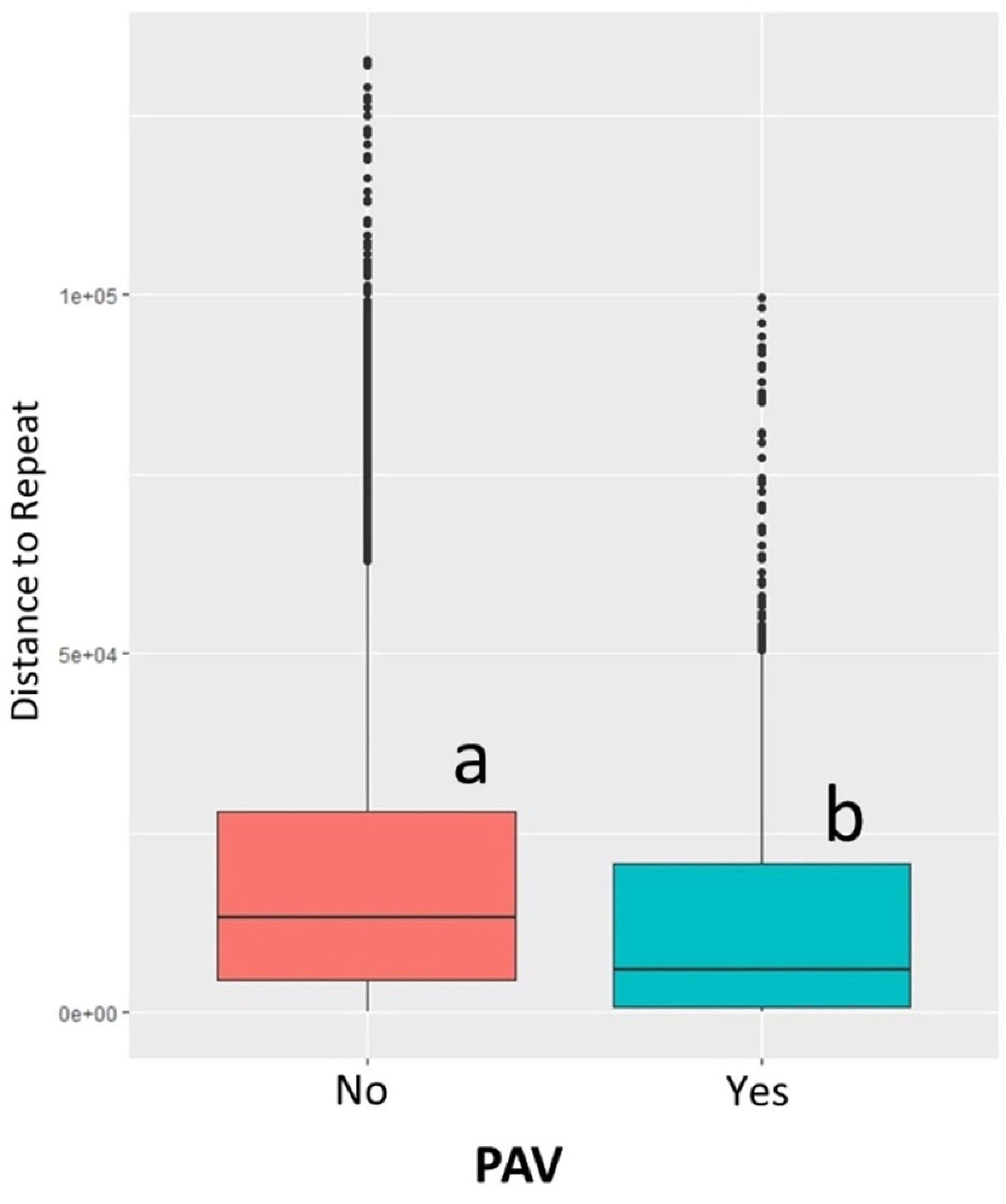
Distances of genes to the nearest repetitive element. Categories are listed on the x-axis and distance to nearest repeat on the y-axis. Letter codes correspond to statistically different groups (Kruskal-Wallis rank sum test p < 0.01).

## Discussion

The dynamic nature of fungal genomes has been revealed following the increase in abundance of whole-genome sequences that facilitate the investigation of the way plant pathogenic fungi undergo diversification. Until recently, effector biology and genome research within *P. nodorum* has relied on relatively few genomic resources. Complementing the recently refined SN15 genome [31] and the establishment of nearly complete telomere to telomere reference genomes of three additional *P. nodorum* isolates [32], this research enhances our knowledge of genomic diversity within *P. nodorum* by revealing locale specific effector diversification and evidence of host susceptibility genes influencing population structure. This investigation also gives insight into the classes of genes undergoing diversifying selection through the accumulation of nonsynonymous SNPs, LOF mutations, and PAV, as well as identifying repetitive elements as a contributing factor to this diversification. Additionally, these results illustrate the utility of these data for the effective implementation of association mapping strategies and the study of variable linkage disequilibrium surrounding effector loci.

### Population structure and genetic diversity

STRUCTURE analysis clearly separated the natural population of *P. nodorum* isolates into two populations. Although isolates collected from North Dakota, Minnesota, and South Dakota were collected from spring, winter, and durum wheat, they still clustered together. Population 2 consisted of isolates from a large geographical range, stretching from Texas to the East coast and further North into Ohio, Maryland, and New York. Overall, a greater number of genic SNPs, both synonymous and nonsynonymous, were observed in Population 1 compared to isolates in Population 2 and is also evidenced in the estimated π values for each population. One possible explanation is the age of the populations. If the United States *P. nodorum* population originated in the Upper Midwest and subsequently spread to the other wheat producing regions, more genome variants may have accumulated. Population size may also play a role in the observed differences and account for the increased level of nucleotide diversity observed in Population 1. Isolates in Population 2 may have recently experienced a bottleneck, resulting in a lower genome-wide Tajima’s D compared to Population 1, as well as the identification of a larger number of potential selective sweep regions. Alternatively, differences may exist in the amount of selection pressure being placed on genes within each population. It has been shown that differences exist in the presence of *P. nodorum* sensitivity genes between wheat germplasm in the Midwest and the Southeastern United States [44, 50]. If the genetic basis of host sensitivity is narrower, with less susceptibility genes present in winter wheat germplasm compared to spring wheat germplasm, selection pressure may have a reduced effect on the genic variation of isolates collected from winter wheat regions.

### Distribution and Diversity of SnToxA, SnTox1, and SnTox3

A large difference was observed in the distribution of *SnToxA* among *P. nodorum* isolates. Only 4.3% of isolates from Population 1 had lost *SnToxA*, compared to the 93.8% of isolates from Population 2, which lacked the gene (Table 2). Additionally, a selective sweep was detected in the genomic region flanking *SnToxA* specifically in Population 1 but was not detected in Population 2 (Fig. 2B). This staggering difference in the prevalence of *SnToxA* is likely correlated with the removal of *Tsn1*, the dominant susceptibility gene that indirectly recognizes SnToxA, from the winter wheat cultivars planted in the soft red winter wheat region of the United States where most of Population 2 was collected. Previous research investigated the sensitivity of winter wheat breeding lines planted in the regions where isolates from the natural population were collected, particularly Georgia, Maryland, and North Carolina. Except for a single line from the breeding program at the USDA-ARS, Raleigh, North Carolina, all breeding lines from the aforementioned regions were found to be insensitive to infiltrations of SnToxA, and therefore, lack a functional *Tsn1* [53]. Conversely, previous research identified SnToxA sensitivity present in popular North Dakota spring wheat cultivars Glenn and Steele ND [54]. Additionally, *Tsn1* is present in the vast majority of wheat cultivars planted in Oklahoma [50]. The strong selective advantage given by SnToxA in the presence of a functional *Tsn1* explains the abundance of isolates harboring this *SnToxA* in Oklahoma and the Upper Midwest. As *Tsn1* is less prevalent in the eastern winter wheat producing regions of the United States, the *SnToxA* gene was lost. A study by McDonald et al. [55] found *SnToxA* to be present in only 25% of a North American *P. nodorum* population, compared to an overall presence of 63.4% in the current natural population. The large difference seen between these two studies is probably due to the differences in the number of isolates used from the Upper Midwest and the rest of the United States. Previously, approximately 8% of the isolates used were from North Dakota, with the remaining isolates being collected from winter wheat producing regions [55], compared to the population used in coverage analysis in this study, consisting of 59.5% of the isolates collected in the Upper Midwest where *Tsn1* is prevalent.

*SnTox1* was the most prevalent *P. nodorum* effector found within this natural population, being present in 95.4% of the isolates (Table 4). The prevalence of *SnTox1* is higher than observed in a North American population by McDonald et al. [55], where it was found to be present in 70% of isolates examined, as well as in 84% of a global population. However, like the study by McDonald et al. [55], *SnTox1* was found to be the most widespread effector of the three characterized genes. Additionally, we detected nine unique haplotypes for *SnTox1* compared to the two private haplotypes previously detected in a North American population [55]. The wide-range distribution of *SnTox1* is likely due to its dual-function in chitin binding and protection from wheat chitinases [56]. The ability of SnTox1 to trigger programmed cell death via recognition by Snn1 is a strong selective force [33, 42] but is dependent on the presence/absence of *Snn1* in locally grown wheat cultivars. The more broad-range effector function of chitinase protection provided by SnTox1, more likely explains its relatively high prevalence compared to the other necrotrophic effectors.

### Gene Diversification

Plant pathogenic fungi are constantly adapting the way they infect their host, often using a suite of effectors to facilitate disease. These effectors are typically small, secreted proteins that lack known functional domains. Additionally, they may be cysteine-rich and exhibit signatures of diversifying selection [57, 58]. Overall, the identified putative effectors in the *P. nodorum* genome exemplify these hallmarks, as evidenced by 72.4% lacking predicted functional domains and accumulating more non-synonymous changes compared to secreted non-effectors or non-secreted proteins. Interestingly, not only are effectors diversifying within the entire population, but effector diversification was also observed to occur within specific subpopulations. As previously discussed, the maintenance of specific host susceptibility genes can shape a population, as seen with the presence of *Tsn1* in hard red spring wheat in the Upper Midwest and the presence of *SnToxA* in nearly every isolate collected in that region, as well as the detection of the genomic region surrounding *SnToxA* as a significant selective sweep specific to Population 1. Selective sweep analysis identified 92 genes underlying sweep regions specific to either subpopulation and no genes located in sweep regions common to both subpopulations. Similar results were observed in the wheat pathogen *Zymoseptoria tritici*, where the investigation of selective sweeps in four globally distinct populations revealed that genomic regions under selection were largely population specific [24]. This highlights the ability of plant pathogenic fungi to rapidly fix advantageous mutations and subsequently shape a population. Additionally, plant pathogenic fungi have developed effector proteins that function in manners other than host colonization, such as competition with local microbial communities [59]. *P. nodorum* likely produces such effectors during its saprotrophic stage and differences in the composition of local microbial populations may be contributing to the diversification of effector proteins within specific pathogen subpopulations. Combined, these results indicate that selection pressure exerted by specific host genes or environmental factors is driving effector/gene diversification as well as fixation and can be restricted to specific geographical regions.

In addition to the diversity observed in predicted effector proteins, gene classes typically associated with pathogen virulence were also observed to exhibit significant levels of polymorphism. Plant chitinases present an obstacle to fungal pathogens via the active degradation of the fungal cell wall and the chitin monomer by-products of this reaction may induce the host defense response [60, 61]. To combat this chitinase activity, fungi have developed chitin binding proteins to protect the fungal cell wall from degradation, as previously observed with *P. nodorum* effector SnTox1 [56]. Another method to counteract host chitinase activity is through the modification of the enzyme by pathogen produced metalloproteases, which reduce chitinase activity resulting in improved virulence [62, 63]. Chitin binding proteins were detected as being enriched within genes exhibiting signs of diversifying selection and LOF mutations in the United States *P. nodorum* population. Additionally, metalloprotease genes were observed to be overrepresented in genes undergoing diversifying selection, as well as containing LOF mutations. This indicates that the interplay between host chitinases and pathogen derived mechanisms of protection may be under strong selective pressure within this population and that the pathogen is not only using a variety of genes to accomplish this, but different means of genic differentiation.

A gene encoding a hydrolase was identified as a candidate virulence gene in a GWAS. Necrotrophs have long been thought to use CWD enzymes for the initiation of infection through the degradation of plant cells [64]. Although necessary for pathogenicity, cell wall degrading enzymes are not typically associated with direct or indirect interactions with host R genes, however, other classes of CWD enzymes have been observed to be under diversifying selection [65, 66]. It is hypothesized that these proteins may trigger the plant basal immune response and are therefore diversifying to avoid this detection [66]. It is also possible that *P. nodorum* hydrolases are evolving rapidly to develop more efficient lysis of plant cells to enhance virulence.

Analysis of repeat content of the genome facilitated the comparison of genomic location of genes and levels of diversity. The clear majority (90.9%) of repetitive DNA identified was classified within families of transposable elements. Genes exhibiting PAV were significantly closer to repetitive elements when compared to genes present in all isolates, suggesting that transposable element activity may contribute to gene gain or loss. This phenomenon has also been observed in *Magnaporthe oryzae* [2] and provides a glimpse into one of the mechanisms giving fungal genomes their plasticity. Additionally, genes residing on the *P. nodorum* accessory chromosome were observed to be evolving faster than genes distributed elsewhere in the genome (Fig. 6). These results further support the hypothesis that transposable element activity mobilizes beneficial genes and upon exposure to significant selection pressure in a given locale, become fixed in a pathogen subpopulation and that different compartments of the fungal genome undergo evolution at different rates.

### Pseudogenization

The detection of polymorphisms within the natural population allowed the analysis of their functional consequences, especially concerning the formation of pseudogenes. Remarkably, a total of 2,848 genes were affected by LOF mutations, amounting to 21.3% of the annotated Sn4 genes. An even greater amount of pseudogenization was observed in the wheat pathogen *Zymoseptoria tritici*, with approximately 55% of the pan-genome harboring at least one LOF mutation [67]. Genes harboring LOF mutations may be functionally redundant and therefore, their losses may have minimal effect on the pathogen. Alternatively, these genes may also be implicated in deleterious interactions with the host, and the pathogen is attempting to escape perception through local adaptation.

### GWAS and LD Decay

Using 322,613 SNP/InDel markers, as well as the PAV of *SnToxA* and *SnTox3* obtained through coverage analysis of whole-genome sequences, a robust framework for association mapping was created. This marker set translates to approximately one marker every 114 bp, overwhelmingly meeting our previous estimate of needing one marker every 7 kb to overcome LD decay [46]. Using four winter wheat lines, the power of GWAS with this dataset was demonstrated, detecting strong associations for the *SnToxA* locus on winter wheat lines Jerry and TAM105, as well as the *SnTox3* locus in winter wheat lines Massey and F/G 95195. Additionally, the utility of this data set was shown by the identification of novel loci associated with virulence on TAM105, enabling the identification of candidate genes for further investigation. In the case of *SnTox3*, the PAV was the most significant marker detected, however, due to the sufficient marker density and LD extending to 6.5 kb, non-causal SNPs were also detected as significant. At the *SnToxA* locus, the PAV of the effector gene was not the most significant variant but was still in high LD with the most significant SNP residing in the region flanking the AT-rich isochore. This is likely due to missing data associated with individual isolates being removed from the PAV dataset due to low coverage across the entire gene set, subsequently reducing the significance of the marker. Interestingly, LD extended much further into the region flanking *SnToxA* compared to that of *SnTox3*. This resulted in the detection of two strong associations in the regions flanking the *SnToxA*-containing isochore. As *SnTox3* is in the subtelomeric region, its flanking region was likely more prone to recombination and the breakdown of LD, as subtelomeric regions are known to be recombination hotspots in fungi [68]. The opposite is likely true at the *SnToxA* locus. Due to the presence/absence nature of the region, recombination may be suppressed and therefore preserve LD.

Prior to this study, focus had been placed on the investigation of diversity within previously identified effector genes. Whole-genome sequencing of 197 *P. nodorum* isolates collected from spring, winter, and durum wheat producing regions of the United States revealed the accumulation of non-synonymous changes in suites of effectors specific to geographical regions. Additionally, *SnToxA* was found to be present in nearly all isolates collected from regions where local wheat lines harbor *Tsn1* and absent from isolates collected from regions where *Tsn1* had been removed from locally grown cultivars. The *SnToxA* locus was detected near a selective sweep region, reinforcing this hypothesis. Selective sweep analysis also revealed unique genomic regions specific to *P. nodorum* subpopulations having undergone purifying selection, indicating that strong and distinct selective forces are acting on each subpopulation. Taken together, this illustrates the selective power that host susceptibility genes place on the pathogen populations and identifies candidate effector genes specific to the spring and winter wheat production regions of the United States. Also, the identified variants were successfully used in a GWAS to identify strong associations of *SnToxA* and *SnTox3* with disease reaction on winter wheat lines, as well as to detect novel virulence candidates. Vastly different patterns of LD were observed surrounding these loci, highlighting the importance of marker density for the successful identification of effector loci in association mapping. Further investigation of *P. nodorum* genomics and functional characterization of effector candidates is ongoing and will provide further information on how this destructive pathogen is interacting with its host.

## Materials and Methods

### DNA Extraction and Whole Genome Sequencing

Dried agarose plugs of each isolate were placed in liquid Fries media [57] and cultured for approximately 48 hours. Fungal tissue was then collected, lyophilized, and homogenized using Lysing Matrix A (MP Biomedicals) by vortexing for 3 minutes on maximum speed. Genomic DNA was extracted using the Biosprint using the manufacturer’s protocol. DNA was enzymatically fragmented using dsDNA fragmentase (New England Biolabs) and whole genome sequencing libraries were prepared using the NEBNext Ultra II Library kit (New England Biolabs) according the recommended protocol. NEBNext Multiplex Oligos for Illumina were used to uniquely index libraries and were subsequently sequenced at the Beijing Genome Institute (BGI) on an Illumina HiSeq 4000. Raw sequencing reads of each isolate were uploaded to the NCBI short read archive under BioProject PRJNA398070.

### Variant Identification

Quality of sequencing reads were analyzed using FastQC [70] and subsequently trimmed using trimmomatic [71]. Trimmed reads were mapped to the *P. nodorum* isolate Sn4 [32] reference genome (NCBI BioProject PRJNA398070) using BWA-MEM [72]. Sequencing coverage of the genome was calculated per isolate using GATK ‘Depth of Coverage’ using default settings. SNPs/InDels were identified using SAMtools ‘mpileup’ [73]. Variants were then filtered for individual genotype quality equal to or greater than 40 with at least three reads supporting the variant and all heterozygous calls were coded as missing data. For GWAS analysis, all SNPs/INDELs with missing data greater than 30% or a minor allele frequency less than 5% were removed from the dataset.

### Population Structure

SNP data were read into the R statistical environment [74] and converted to a genlight object using the package ‘vcfR’ [75]. PCA was conducted within the R package ‘adegenet’ [76] using a randomly selected subset of 50,000 markers.

A marker subset minimizing potential linkage disequilibrium between marker pairs consisting of approximately one SNP/10 kb was created in TASSEL v5 [77] and used to infer population structure using the software STRUCTURE [78]. A burn-in of 10,000 followed by 25,000 MCMC replications using the admixture model was completed for each k value from 1-8 with three iterations. Optimal sub-population level was determined using the method developed by Evanno et al. [79] and implemented in StructureHarvester [80]. STRUCTURE analysis was re-run using the optimal k value with a burn-in period of 10,000 and 100,000 MCMC. An individual isolate was assigned to a specific subpopulation if membership probability was greater than 0.85. Subsets corresponding to SNP/InDel calls for individuals within specific populations were created using VCFtools [81]. ADMIXTOOLS [49] was used to conduct a 3-population test to determine if evidence existed for the hypothesis that isolates collected from Oklahoma resulted from admixture between Population 1 (Upper Midwest) and Population 2 (South/East United States). Genotypic data was converted to PLINK format in TASSEL 5. Genotypic data was further converted to EIGENSTRAT format using ‘convertf’ provided in ADMIXTOOLS. A 3-population test was conducted using the two major populations identified using STRUCTURE as the sources and the isolates collected from Oklahoma as the target population. As recommended in the software documentation, the ‘inbreed’ option was selected due to the haploid nature of the organism. Additionally, as the f_3 statistic uses a measure of heterozygosity for normalization, as recommended in the documentation, the parameter ‘OUTGROUP=YES’ was used. Isolates collected from Oklahoma (n=17), not clearly belonging to either major population, were removed from the datasets when comparing pN/pS ratios to obtain a more representative estimate of diversifying or purifying selection for each population.

### Population Genomics

Using genotypic data for 197 *P. nodorum* isolates and population assignments derived from STRUCTURE analysis, population genomics analyses were conducted in the R package ‘PopGenome’ specifying a ploidy level of one [82]. F_st_, nucleotide diversity (π), Watterson’s Θ, and Tajima’s D were calculated across individual chromosomes and averaged to obtain a genome-wide value. Tajima’s D was also calculated in 50 kb windows in 25 kb steps across each chromosome to examine regions that substantially lack allelic diversity and may provide evidence for selective sweeps.

A likelihood-based method implemented in SweeD [83] was used to detect regions of the genome that have undergone a selective sweep. Chromosomes were divided into approximate 1 kb grids and each population was analyzed separately. The option ‘-folded’ was selected to consider the site frequency spectrum as folded, as to not distinguish between ancestral and derived states. Grids within the 99^th^ percentile of likelihood values were extracted and examined for values of Tajima’s D. If the value of Tajima’s D fell below zero within an interval, it was considered as a selective sweep region. Selective sweep grids within 10 kb were considered a single region. Genes underlying selective sweep regions were extracted using BEDtools ‘intersect’ [84] and gene ontology enrichment analysis was conducted as described in a subsequent section.

### Disease Phenotyping and Effector Infiltration

The natural *P. nodorum* population (n=197) used in this study consists of 51 isolates collected from spring wheat in North Dakota and Minnesota, 45 isolates collected from durum wheat in North Dakota, nine isolates collected from winter wheat in South Dakota, and 92 isolates collected from winter wheat in the Eastern, Southern, and Pacific Northwest regions of the United States (S1 Table). These isolates were chosen for their diversity in geographical origin of isolation, as well as the differences in wheat class (spring, winter, and durum wheat). *P. nodorum* isolates were cultured as described by Friesen and Faris [69]. Briefly, dried agarose plugs stored at −20 °C of each isolate were brought to room temperature and rehydrated by placing on V8-potato dextrose agar (150 mL V8 juice, 3 g CaCO_3_, 10 g Difco PDA, 10 g agar, 850 mL H_2_O). Plugs were then spread across the plate to distribute the spores and the plates were then incubated at room temperature under a constant fluorescent light regimen for seven days. Pycnidial spores were harvested by flooding the plates with sterile H_2_O and agitation with a sterile inoculation loop. Spores were counted using a hemocytometer, concentration was adjusted to 1 × 10^6^ spores per mL, and two drops of Tween20 were added per 100 mL of spore suspension.

Disease reaction to the 197 *P. nodorum* isolates was assessed on six wheat lines including spring (Alsen) and winter types (Jerry, TAM105, Massey, and F/G95195), as well as a recombinant from a synthetic wheat population (ITMI38). Three seeds of each wheat line were sown into cones, comprising a single replicate. A border of wheat cultivar Alsen was planted to reduce the edge effect. Plants were grown in the greenhouse for approximately two weeks, until the two to three-leaf stage. Inoculations were conducted as described by Friesen and Faris [69]. Briefly, plants were inoculated using a paint sprayer until leaves were fully covered, until runoff. Inoculated plants were placed in a mist chamber at 100% humidity for 24 hours and subsequently moved to a climate-controlled growth chamber at 21 °C with a 12 hour photoperiod for six days. Disease ratings were taken seven days post-inoculation using the 0-5 scale as described by Liu et al. [85]. For each wheat line, at least three replicates were conducted per *P. nodorum* isolate and the average of the replicates were used as the phenotypic data for downstream analyses.

To produce effector proteins used in bio-assays to validate associations detected with known necrotrophic effectors, cells from glycerol stocks of *P. pastoris* expressing previously characterized SnToxA and SnTox3 were plated on YPD media amended with Zeocin (Invitrogen) at 25 µg/mL and incubated at 30 °C for 2-4 days until colony formation [35, 36]. Colonies were picked and grown in 2 mL of liquid YPD media and grown for 24 hours at 30 °C and shaking at 250 rpm. A 500 µL aliquot of each starter culture was transferred to 50 mL of YPD media and incubated at 30 °C and shaking at 250 rpm for 48 hours. Cultures were transferred to 50 mL conical tubes and centrifuged at 3,166 rcf for 10 minutes. Supernatants were decanted and filtered using a 0.45 µm Durapore Membrane Filter (Merck Millipore Ltd.). Filtered supernatant was then lyophilized overnight and stored at −20 °C. Seeds of winter wheat lines Jerry, TAM105, Massey, and F/G95195 were sown in cones and grown under greenhouse conditions for approximately 14 days until the secondary leaf was fully emerged. Freeze-dried protein samples of SnToxA and SnTox3 were resuspended in ddH_2_O and infiltrated into secondary wheat leaves using a needleless syringe. Plants were placed in a growth chamber at 21 °C with a 12-hour photoperiod and evaluated for necrotrophic effector sensitivity after three days.

### Genome-Wide Association Analysis

Association mapping was conducted using TASSEL v5 [77] and GAPIT [86, 87]. A naïve model, as well as a model including the first three components derived from a PCA as fixed effects were used in association analyses in TASSEL v5. A kinship matrix (K) was calculated using the EMMA algorithm. Models incorporating K as a random effect, as well as a model using a combination of PCA and K were analyzed in GAPIT. The most robust model was chosen for each trait by visualization of Q-Q plots produced by GAPIT, illustrating the observed vs expected unadjusted p-values. Marker p-values were adjusted using a false discovery rate (FDR) in the R Statistical Environment. Markers with a FDR adjusted p-value of 0.05 or less were deemed significant.

### Linkage Disequilibrium

Linkage disequilibrium (LD) was calculated between the most significant marker detected from association mapping analyses and each intrachromosomal marker in TASSEL v5 [77]. LD decay was determined by averaging R^2^ values across sliding windows consisting of five markers and observing when average R^2^ values fell below 0.20 for three consecutive windows. Position of the extent of LD was taken as the position of the marker at the center of the sliding window before decay.

### Putative Effector Diversification Analyses

The presence of predicted secretion signals and transmembrane domains were identified using DeepSig [88]. Effector candidates were predicted via EffectorP using protein sequences of secreted proteins [89]. For comparative analyses, proteins were categorized as effectors (identified by EffectorP), secreted non-effectors (predicted secreted proteins without transmembrane domain or effector predictions), or non-secreted. Genomic positions of all genes were obtained and used to calculate genome coverage with BEDtools ‘coverage’ [84]. A gene was classified as absent if sequencing coverage across the entire length of the gene was less than 40%.

A consensus genome sequence was obtained for each isolate using bcftools consensus, specifying “--sample” [90]. The perl script ‘gff2fasta.pl’ was used to extract the coding regions for each isolate separately (https://github.com/minillinim/gff2fasta/blob/master/gff2fasta.pl). Coding regions of each gene were aligned using Clustal Omega [91] and the subsequent alignments were concatenated. The average number of nonsynonymous (NSite) and synonymous sites (SSite) were calculated using egglib [92].

Synonymous (S) and non-synonymous (N) SNPs, as well as LOF mutations were identified using SNPeff [93] and SNPsift [94] with the *P. nodorum* isolate Sn4 reference genome annotation. LOF mutations include small InDels causing frameshifts, as well as SNPs/InDels resulting in the loss of a start codon, loss of a stop codon, or gain of a premature stop codon. Isolates from specific subpopulations were grouped and fixed SNPs/LOF mutations (minor allele frequency of 0%) were removed from each dataset. pN/pS ratios were obtained for each gene by the following formula: (N/NSites)/(S/SSites). Genes with pN/pS ratios greater than one were considered to be under diversifying selection, while genes with pN/pS ratios equal to or less than one were considered to be under neutral or purifying selection. Raw counts of LOF mutations were used in comparisons between subpopulations. Protein sequences of the *P. nodorum* isolate Sn4 annotation were input into InterproScan [95] to identify conserved domains and gene ontology. Gene ontology enrichment analysis was conducted in the R Statistical Environment using the package ‘topGO’ [96]. Analysis was conducted on both molecular function and biological process associated ontology terms using the classic algorithm and significance of overrepresented terms were determined using Fisher’s exact test. GO terms with FDR adjusted p-values of less than 0.10 were declared significantly enriched.

### Repetitive Element Annotation and Analysis

De novo identification of repetitive element families in the Sn4 reference genome was conducted using RepeatModeler [97]. Identified repetitive element families were used as input to RepeatMasker [98] to annotate the repetitive regions of the Sn4 genome. Distance to the nearest repetitive element from each annotated gene was calculated using bedtools ‘closest’ [84].

## Data availability

All novel data used in analyses described within the manuscript are available at https://github.com/jkzrich/pnodorum_popgen.

## Acknowledgements

The authors would like to thank Danielle Holmes and Jason Axtman for technical assistance. The project was funded by the United States Department of Agriculture (USDA) National Institute of Food and Agriculture, Agriculture and Food Research Initiative Competitive Grant number 2016-67013-24813. Mention of trade names or commercial products in this publication is solely for the purpose of providing specific information and does not imply recommendation or endorsement by the USDA. USDA is an equal opportunity provider and employer.

---

**S1 Figure.** STRUCTURE analysis of 197 *P. nodorum* isolates. A) Evanno method indicated the optimal k value (number of subpopulations) to be two. B) Using a k-value of two, the 197 *P. nodorum* natural population is divided into two populations. Population 1 (red) consists of isolates from the Midwestern United States. Population 2 (blue) consists of isolates from the Southern/Eastern region of the United States, as well as Oregon.

**S2 Figure.** Boxplots illustrating the distribution of (A) nucleotide diversity, (B) Tajima’s D, and (C) Watterson’s Theta calculated in 50 kb windows within Population 1 and Population 2.

**S3 Figure.** Whole genome Manhattan plots illustrating significant associations with virulence on wheat lines TAM105, Jerry, Massey, and F/G95195. Dots represent individual SNPs/InDels. Markers are ordered by position and chromosomes are displayed on the x-axis. The –log10(p) value is displayed on the y-axis. The horizontal line represents the significance threshold at and FDR adjusted p-value of 0.05.

**S1 File.** Results of the gene ontology enrichment analysis conducted using topGO.

**S2 File.** Phenotypic data

**S1 Table.** Collection information of the fungal isolates used in this study.

**S2 Table.** Genome sequencing coverage statistics for each isolate.

**S3 Table.** Genes under population-specific diversifying selection.

**S4 Table.** STRUCTURE likelihoods

